# Distinct neural input strategies to motor units in the soleus and medial gastrocnemius during quiet standing

**DOI:** 10.64898/2026.01.11.698550

**Authors:** Hiroki Arakawa, Naotsugu Kaneko, Nadaka Hakariya, Hikaru Yokoyama, Kimitaka Nakazawa

**Author notes:** authors are co-corresponding authors., Corresponding author H. Yokoyama: Institute of Engineering, Tokyo University of Agriculture and Technology, Tokyo, Japan., K. Nakazawa: Department of Life Sciences, Graduate School of Arts and Sciences, The University of Tokyo, Tokyo, Japan.

## Abstract

Standing posture control critically depends on the activation of the soleus (SOL) and medial gastrocnemius (MG), which serve distinct functional roles. Identifying underlying neural mechanisms has been challenging, as conventional invasive techniques sample only limited motor units (MUs). Recent advances in high-density surface electromyography (HDsEMG) have enabled analysis of MU activity and estimation of common synaptic inputs to spinal motoneurons. Therefore, we aimed to elucidate the common synaptic inputs underlying the distinct MU behaviors of SOL and MG during standing. We recorded HDsEMG from the SOL and MG, alongside electroencephalography, from 20 male participants during standing and isometric voluntary contractions. EMG signals were decomposed into individual MU activity, with common synaptic inputs estimated through intramuscular and corticomuscular coherence analyses (IMC and CMC). Compared to SOL, the MG exhibited significantly higher delta-, alpha-, and beta-band IMC during standing. In task comparisons, alpha-band IMC increased during standing specifically in the MG. Furthermore, although beta-band CMC decreased in both muscles while standing, IMC was preserved in the MG but markedly reduced in SOL. This dissociation suggests that the common neural drive to the MG during standing is likely derived from subcortical rather than cortical pathways. These results demonstrate that the SOL and MG are governed by distinct neural control strategies, which likely underlie their functional roles. Given the low CMC, the MG relies on strong common synaptic input from subcortical pathways (e.g., vestibulospinal and reticulospinal) to produce rapid corrective torque, whereas the SOL functions with lower neural synchrony to ensure steady ankle stiffness.

**Key points:**

- The soleus and medial gastrocnemius play distinct roles in standing control, however, due to technical limitations, it has been difficult to identify the underlying neural mechanisms responsible for these differences.
- Using high-density surface electromyography, we examined motor unit activity and neural inputs to these muscles during standing.
- The medial gastrocnemius shows greater common synaptic input, potentially facilitating rapid ankle plantarflexion torque generation to correct postural sway.
- The soleus exhibits lower motor unit synchrony, enabling stable and continuous ankle plantarflexion torque generation for body weight support.
- This study demonstrates that the soleus and medial gastrocnemius are governed by distinct neural control strategies, which likely underlie their distinct functional roles.

**Abstract figure legend:** 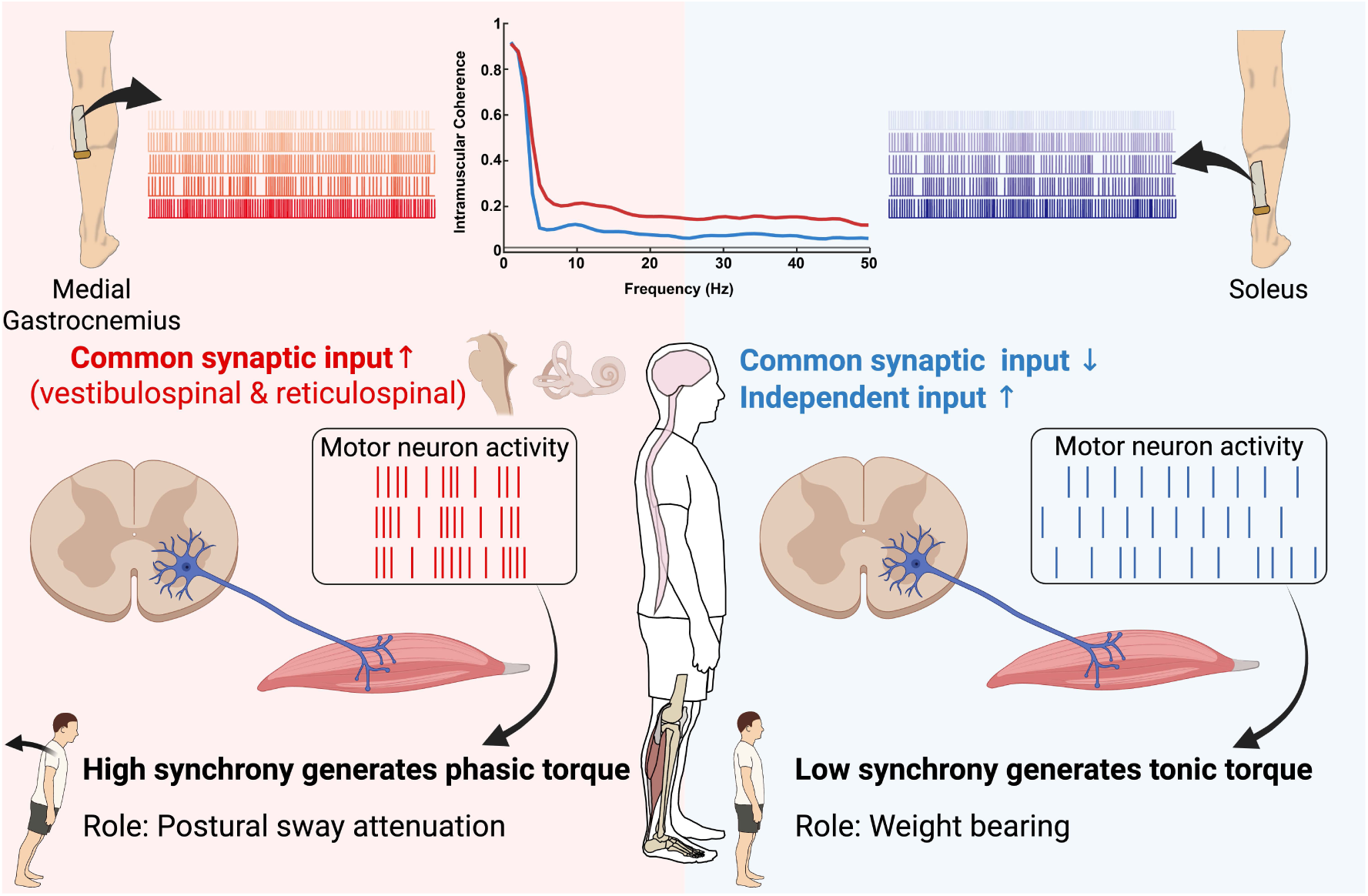

Motor unit spike trains were decomposed from high-density surface electromyograms recorded from the medial gastrocnemius (MG; left; red) muscle and soleus (SOL; right; blue). To compare the neural input to the spinal motor neurons between them, we quantified the intramuscular coherence (IMC) of motor unit spike trains within the delta, alpha, and beta bands. MG exhibited greater IMC than SOL during standing, indicating stronger common synaptic input, likely mediated by vestibulospinal and reticulospinal pathways. In contrast, SOL showed lower IMC, suggesting a greater contribution of independent synaptic input. As a consequence, high motor-unit synchrony in the MG supports rapid, phasic torque generation for postural sway attenuation, whereas low synchrony in the SOL enables smooth, steady torque production for weight bearing during standing.

## Introduction

Standing posture control represents a fundamental aspect of human motor control. During quiet standing, the body’s center of mass is positioned slightly anterior to the ankle joint, requiring continuous ankle plantarflexion torque to prevent falling forward (Loram & Lakie, 2002; Winter, 2009). Among the triceps surae, the soleus (SOL) and medial gastrocnemius (MG) are the primary muscles responsible for producing this torque, with a smaller contribution from the lateral gastrocnemius (Loram *et al*., 2005; Héroux *et al*., 2014). Although SOL and MG play essential roles, their functions during quiet standing differ (Héroux *et al*., 2014). The SOL exhibits relatively stable, tonic activity, providing sustained ankle plantarflexion torque to support body weight (Masani *et al*., 2003; Héroux *et al*., 2014). In contrast, the MG shows large fluctuations that are synchronized with center-of-mass sway, thereby contributing to the attenuation of postural sway (Masani *et al*., 2003; Héroux *et al*., 2014). These functional differences are often attributed to anatomical factors. The MG, a biarticular muscle with a high proportion of fast-twitch fibers, is suited for rapid, phasic force production, whereas the SOL, a monoarticular muscle with predominantly slow-twitch fibers, is specialized for sustained, tonic force generation (Buchthal & Schmalbruch, 1980). However, these structural characteristics do not fully account for the pronounced differences in their activation during standing. This suggests that distinct neural inputs may also underlie the functional specialization of the SOL and MG.

In terms of neural factors, previous studies have demonstrated that the SOL and MG differ in their responsiveness to spinal and descending inputs during quiet standing. The SOL generally shows a greater H-reflex amplitude than the MG (Hayashi *et al*., 1992; Makihara *et al*., 2012), likely reflecting its predominance of slow-twitch fibers and the associated lower motor neuron recruitment thresholds (Buchthal & Schmalbruch, 1980). In contrast, responses to vestibular stimulation are larger in the MG than in the SOL (Dakin *et al*., 2016), suggesting that vestibular inputs exert a stronger influence on MG motor neurons. These observations indicate that sensory inputs, including Ia afferent and vestibulospinal pathways, help shape the distinct behavior of the SOL and MG during standing. However, these structural characteristics alone are insufficient to explain the magnitude and task specificity of the observed differences, implying that additional pathways, such as reticulospinal and corticospinal inputs, also contribute to their neural control.

To clarify how neural inputs contribute to muscle-specific roles in postural control, it is necessary to examine how these inputs are expressed at the level of spinal motor neurons. Because motor neuron output is expressed through the firing patterns of its motor units (MUs), MU activity most directly reflects the integrated neural drive received by each muscle. Thus, differences in MU firing behavior and synchrony offer critical insights into how distinct neural inputs shape the functional roles of the SOL and MG during standing. Traditionally, investigations of MU activity have relied on intramuscular needle or wire electrodes, which are invasive and restricted to a limited number of identifiable MUs. This limitation has long prevented a comprehensive understanding of the neural inputs to the spinal motor neuron pool. In recent years, however, the advent of high-density surface electromyography (HDsEMG) has transformed this field. HDsEMG enables the non-invasive identification of a large population of MU activities, thereby overcoming the constraints of conventional invasive techniques. This technique provides direct access to the integrated neural inputs to spinal motor neurons, thereby offering an approach to investigate muscle-specific neural control strategies during quiet standing.

By leveraging this methodology, we aimed to elucidate the neural control mechanisms underlying the distinct functional roles of the SOL and MG during quiet standing. To this end, we employed coherence analysis, a method that evaluates frequency-specific correlations between two signals to infer the neural inputs to spinal motor neurons (Cadzow & Solomon, 1987; Farmer *et al*., 2007). Intramuscular coherence (IMC)—measured as the coherence between two cumulative spike trains (CSTs) created by summing the activity of different sub-groups of motoneurons—serves as an indicator of the common synaptic input to spinal motoneurons (Castronovo *et al*., 2015). By simultaneously examining corticomuscular coherence (CMC) between the CST and activity in the sensorimotor cortex, which reflects functional connectivity between the cortex and spinal motoneurons (Conway *et al*., 1995), we can infer how strongly cortical drive contributes to this common input. Importantly, coherence in different frequency bands is thought to reflect contributions from distinct neural pathways. By integrating these approaches with HDsEMG, this study provides novel evidence on how neural inputs modulate MU activity in the SOL and MG during standing, offering new insights into the neural control of upright posture.

## Method

### Participants

This study included 20 male participants (height, 173.2 ± 5.0 cm; weight, 68.0 ± 4.9 kg; age, 25.1 ± 2.7 yr) with no known neuromuscular disorders. Informed written consent was obtained from all participants prior to the start of the experiment. Since females generally yield fewer identifiable MUs during HDsEMG decomposition, likely due to factors such as thicker subcutaneous tissue (Taylor *et al*., 2022; Jenz *et al*., 2023), only male participants were included in the present study in line with prior work on the HDsEMG decomposition (Del Vecchio *et al*., 2019; Martinez-Valdes *et al*., 2020). The current study was approved by the Ethics Committee of the University of Tokyo (approval number: 873-3)

### Experimental Procedure

Figure 1 shows an overview of the experimental design. Participants performed two tasks: a quiet standing task and an isometric voluntary contraction task. For the quiet standing task, participants stood quietly with their eyes open for 2 minutes, maintaining a relaxed upright posture while focusing on a visual target placed in front of them (Fig. 1A). For the isometric voluntary contraction task, participants were seated with the knee joint fully extended and the ankle joint fixed at 0° using an ankle–foot orthosis (Fig. 1A). In this task, they were instructed to match their muscle activity to a target displayed on a monitor. The target corresponded to the mean EMG level of the MG or SOL recorded during the standing task, and participants were asked to reproduce this level of EMG as accurately as possible. The tasks were performed separately for the MG and SOL, with each muscle tested twice (two 30-second trials per muscle).

**Figure 1.**
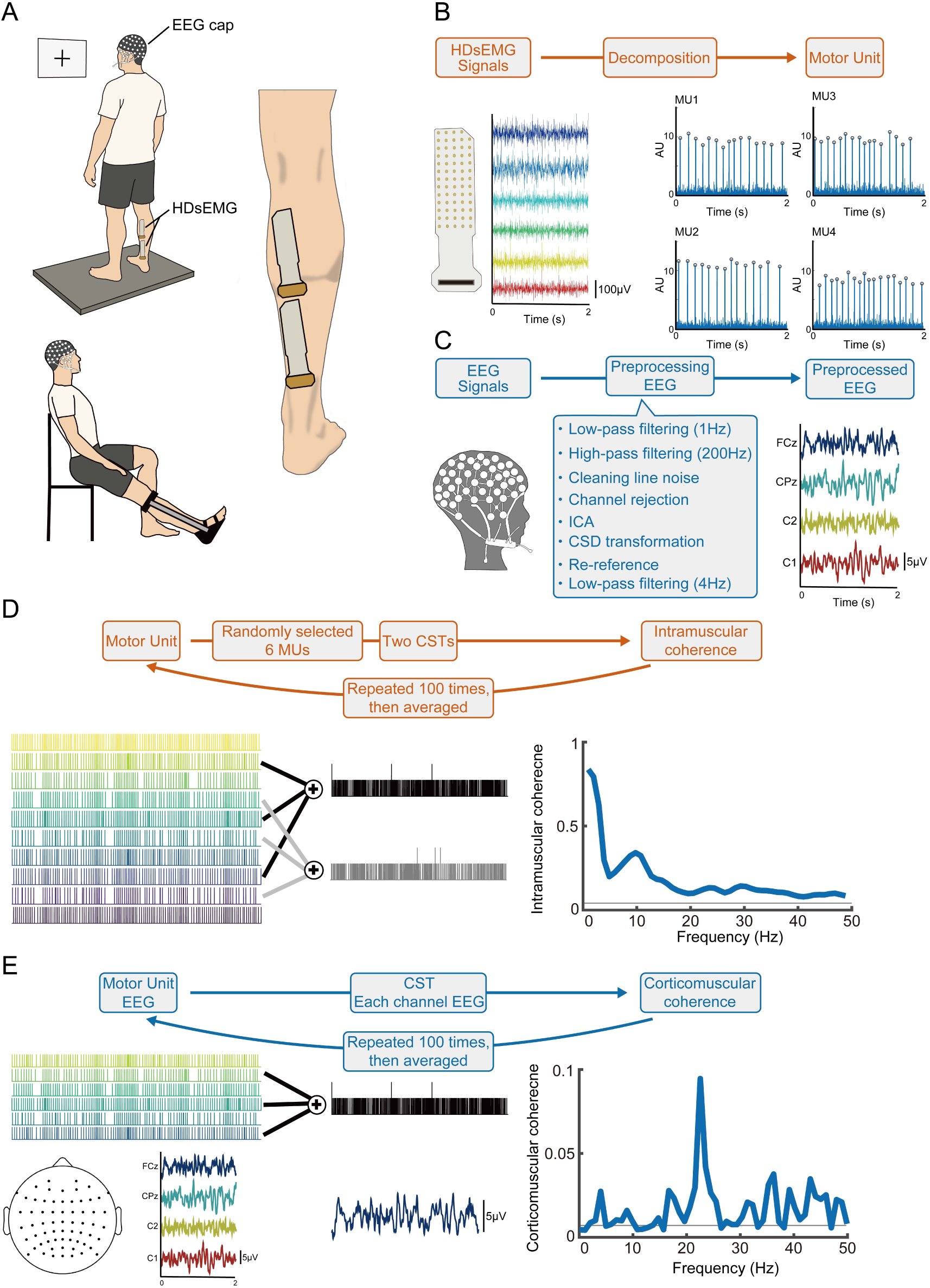
Experimental setup and Method for common synaptic input estimation. **A**. schematic representation of the participants’ posture for measuring muscle activity during standing and isometric voluntary contraction. High-density surface electromyograms (HDsEMG) were acquired from the medial gastrocnemius (MG) and soleus (SOL) using a grid of 64 electrodes. HDsEMG and electroencephalography (EEG) were recorded simultaneously. **B**. Schematic illustration of motor unit (MU) identification. HDsEMG signals recorded from the muscle were decomposed to extract the discharge timings of individual MUs. **C**. Schematic representation of EEG preprocessing. Raw EEG signals were preprocessed. **D**. Analysis workflow for calculating IMC. **E**. Analysis workflow for calculating CMC.

**Figure 2.**
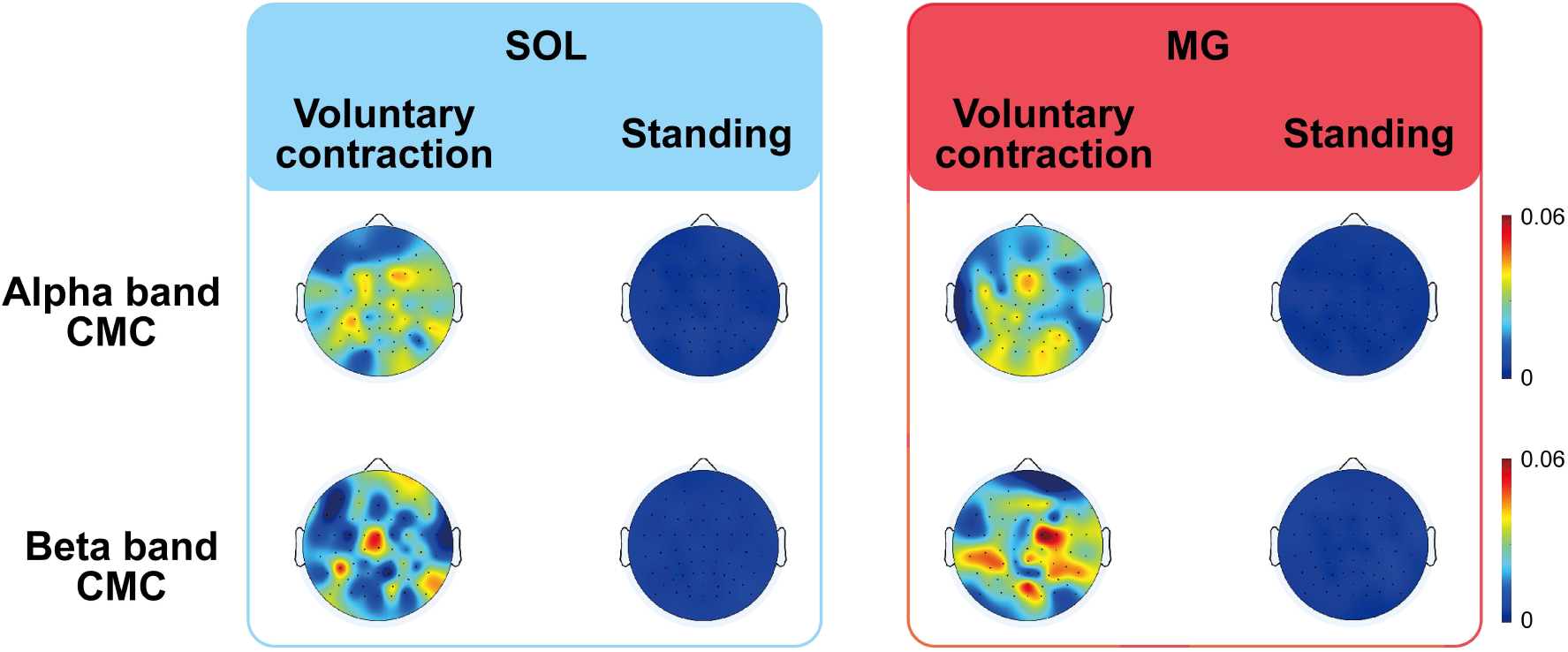
Spatial distribution of corticomuscular coherence across the scalp. The top four topographies represent CMC in the alpha band, whereas the bottom four represent CMC in the beta band.

### Data Collection

HDsEMG signals were recorded from the right MG and SOL using a two-dimensional adhesive grid of 13 × 5 equally spaced electrodes (each 1 mm in diameter, inter-electrode distance 8 mm), with one electrode absent from the upper right corner (GR08MM1305, OT Bioelettronica, Torino, Italy) (Fig.1A). The skin over the MG and SOL was shaved and cleansed with abrasive gel and alcohol wipes to minimize electrode–skin impedance. Electrode-skin impedance was reduced using conductive paste (Ten 20 conductive paste, Weaver and Company, Colorado, USA). HDsEMG signals were sampled at 2048 Hz and band-pass filtered between 10 and 500 Hz with a bio-amplifier (Quattrocento, OTBioelettronica, Torino, Italy). Prior to the HDsEMG decomposition analysis, 64 monopolar EMG signals were re-referenced to generate 59 bipolar signals by calculating the difference between signals from adjacent electrodes aligned along the longitudinal axis of the muscle.

Electroencephalography (EEG) was recorded using a sponge-based passive electrode cap (R-Net, Brain Products, Gilching, Germany) with an EEG amplifier (actiCHamp, Brain Products, Gilching, Germany), sampled at 1000 Hz (Fig.1A). The electrodes positioned according to the international 10-10 system. Electrode impedances were maintained below 30 kΩ, which was substantially lower than the recommended threshold of 50 kΩ for high-impedance EEG amplifiers.

### Data analysis

#### EMG low-frequency components

For the standing task, a 1-minute segment, beginning 30 seconds after the trial start, was used for analysis. For the isometric voluntary contraction task, two 30-second trials were concatenated to obtain a 1-minute analysis interval. For each task, the 59-channel differential HDsEMG signals were rectified. After that, they were low-pass filtered at 4 Hz using a fourth-order, zero-phase-lag Butterworth filter to extract the low-frequency components of the EMG (Masani *et al*., 2003; Winter, 2009). To quantify the variability of these low-frequency components, the coefficient of variation (CoV) was calculated for each channel, and the mean value across all channels was then obtained.

#### HDsEMG decomposition

HDsEMG signals were decomposed into individual MU spike trains using an automatic convolutive blind source separation (BSS) algorithm, which combines fast independent component analysis with the convolutional kernel compensation method (Negro *et al*., 2016) (Fig.1B). Decomposition was performed separately for the HDsEMG recordings obtained during the standing and isometric voluntary contraction tasks. The quality of the decomposition was assessed using the silhouette (SIL) value, which quantifies the clustering quality of extracted discharge timings. Specifically, SIL is calculated as the difference between the within-cluster and between-cluster sums of point-to-centroid distances, normalized by the larger of the two values (Negro *et al*., 2016). In this study, only MUs with a SIL value greater than 0.88 were retained for further analysis (Negro *et al*., 2016; Cabral *et al*., 2024*a*; Gomes *et al*., 2024). For individual MUs, we calculated the firing rate and the number of recruited MUs for each task. For the firing rate during standing, inter-spike intervals (ISIs) longer than 250ms were excluded, as these reflect silent periods caused by postural sway-induced modulation (Héroux *et al*., 2014). In addition, ISIs shorter than 5ms were also omitted to avoid physiologically implausible firing rates.

#### EEG pre-processing

EEG analysis was conducted using custom scripts incorporating EEGLAB v2024.2 functions (Fig.1B). The EEG signals were high-pass filtered at 1 Hz and low-pass filtered at 200 Hz using the “eegfiltnew”. Power line noise (50 and 100 Hz) was removed using the “Cleanline” function. Noisy channels were removed using the pop-rejchan function. Channel exclusion was based on kurtosis (SD = 5) and probability (SD = 10) criteria. The EEG data were subsequently re-referenced to the common average. Following preprocessing, the EEG data were decomposed using Adaptive mixture independent component analysis (AMICA) for artifact removal (Studnicki *et al*., 2023). The resulting independent components (ICs) were automatically classified and labeled using the pop_iclabel function, which assigns probabilistic labels to each IC across seven categories: brain, muscle, eye, heart, line noise, channel noise, and other.

Components labeled as “brain” or “other,” as well as any components whose probability for the assigned category was below 75%, were retained for further analysis (Studnicki *et al*., 2023). These ICs were projected back onto the channel, resulting in the retention of 62.2 ± 1.14 channels. This procedure effectively reduced artifacts such as eye blinks, muscle activity, and cardiac signals, while preserving brain-related activity. To enhance spatial resolution and minimize volume conduction effects, the current source density (CSD) transformation was applied to the cleaned EEG data (Kayser & Tenke, 2006). The CSD was computed using the spherical spline method implemented in the CSD toolbox, with the standard 10–10 electrode configuration and default parameters.

#### Power spectrum density (PSD) analysis

We analyzed the PSD of surface EMG and EEG. PSDs were computed using Welch’s method with a 1-s segment length and 50% overlap.

For the surface EMG PSD, rectified EMG signals from the proximal (channel 25), central (channel 29), and distal (channel 33) electrodes of the HDsEMG grids placed over the MG and SOL were used. PSD was normalized by the total PSD and expressed as a percentage (%PSD). The PSDs from these channels were averaged to represent the overall spectral characteristics of each muscle. The mean %PSD within the delta (1–5 Hz), alpha (5–15 Hz), and beta (15–30 Hz) frequency bands was then calculated (Cabral *et al*., 2024*b*; Contreras-Hernandez *et al*., 2025).

For the PSD of EEG, signals were obtained from electrodes covering the sensorimotor region (Cz, C1, C2, C3, C4, FCz, FC1, FC2, FC3, FC4, CPz, CP1, CP2, CP3, CP4). PSD was then computed for each channel, and the mean PSD across these channels was used (Nazarpour *et al*., 2006; Edelman *et al*., 2016). We quantified the average power within the alpha (8–13 Hz), beta (13–30 Hz), and gamma (30–40 Hz) frequency bands (Pernet *et al*., 2020).

#### Intramuscular coherence (IMC) analysis

We evaluated MU synchronization in the frequency domain among the MU spike trains from the same muscle to compare common synaptic input to motor neurons between standing and voluntary contraction tasks. We performed coherence analysis on two cumulative spike trains (CSTs) of equal size (Fig.1C). In this study, periods in the CSTs with no discharges longer than 250 ms were considered silent periods caused by postural sway-induced modulation and were excluded from the analysis. Each CST was constructed by summing the binary discharge trains of three MUs (Rossato *et al*., 2022; Yokoyama *et al*., 2022). For each analysis iteration, six MUs were randomly selected from the pool of identified units—three for each CST. This random grouping process was repeated 100 times to ensure robustness of the coherence estimates. Coherence was calculated as follows:

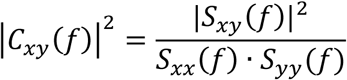

where S_xy_(f) is the cross-spectrum between x and y, and S_xx_(f) and S_yy_(f) are the auto spectra for x and y, respectively, for frequency f. The coherence was computed in 400 ms nonoverlapping windows. The following equation defines the significance level of coherence:

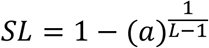

where α is the confidence level (0.05), and L is the number of segments (steps) used for the analysis. In this study, we calculated the coherence area within the delta (1–5 Hz), alpha (5–15 Hz), and beta (15–30 Hz) frequency bands (Cabral *et al*., 2024*b*; Contreras-Hernandez *et al*., 2025).

#### Corticomuscular coherence (CMC) analysis

CMC was calculated using the same equations described above for the comparison between EEG and CST data (Fig.1C). Initially, CMC was computed between EEG signals recorded from 64 scalp electrodes and EMG signals from the right medial gastrocnemius (MG) and soleus (SOL) muscles to determine the scalp distribution of CMC. For statistical analyses, the maximum CMC value among 64 electrodes was used. Consistent with IMC analyses, we calculated the coherence area within the delta (1–5 Hz), alpha (5–15 Hz), and beta (15–30 Hz) frequency bands (Cabral *et al*., 2024*b*; Contreras-Hernandez *et al*., 2025).

Five participants were excluded from the coherence analyses (IMC and CMC) because fewer than six MUs were identified in either the SOL or MG. Thus, data from 15 participants were used for these analyses.

### Statistical analysis

Prior to conducting parametric analyses, the normality of the within-subject differences for each condition was assessed using the Shapiro–Wilk test. The mean EMG amplitude, the number of MUs, and the mean firing rate were confirmed to be normally distributed; therefore, paired t-tests were performed to compare the mean EMG amplitude, the number of MUs, and the mean firing rate across tasks. On the other hand, the CoV of low-frequency EMG was not normally distributed; thus, the Wilcoxon signed-rank test was performed to compare the CoV between tasks and between muscles.

Since the EEG PSD data were not normally distributed, the Friedman test was used to compare the PSD of alpha (5–13 Hz), beta (13–30 Hz), and gamma (30–40 Hz) bands across the three conditions (standing, SOL isometric voluntary contraction, MG isometric voluntary contraction). Kendall’s W was calculated to estimate the effect size. When a significant main effect was observed, post-hoc pairwise comparisons were conducted using Wilcoxon signed-rank tests with Bonferroni correction. For these pairwise comparisons, the effect size r was calculated as Z /√N. For both Kendall’s W and effect size r, a value less than 0.1 was classified as ‘trivial’, 0.1–0.3 as ‘small’, 0.3–0.5 as ‘moderate’, and greater than 0.5 as ‘large’ <u>(Cohen, 2009)</u>.

A two-way repeated-measures ANOVA was performed for both the coherence measures (IMC and CMC) and the PSD of the surface EMG. For each frequency band (delta: 1–5 Hz, alpha: 5–15 Hz, beta: 15–30 Hz), posture (sitting vs standing) and muscle (SOL vs. MG) were included as within-subject factors, with the subject treated as a random effect. For significant effects or interactions, post hoc paired t-tests were performed, and Bonferroni correction was applied for multiple comparisons (4 comparisons per band). Adjusted p-values were reported. All statistical analyses were performed using R (version 4.4.3), with significance set at p < 0.05. For a two-way repeated-measures ANOVA, effect sizes were reported as eta squared (η²) values. For post hoc paired t-tests, Cohen’s d was calculated as a measure of effect size. A Cohen’s d less than 0.2 was classified as ‘trivial’, 0.2–0.5 as ‘small’, 0.5–0.8 as ‘moderate’, and greater than 0.8 as ‘large’ (Cohen, 2009).

## Results

### EMG and MU firing behavior

There were no significant differences between standing and isometric voluntary contraction in mean EMG amplitude, number of motor units, or firing rates for either the SOL or the MG (Fig. 3A–C). Specifically, mean EMG amplitude was comparable between tasks in both muscles (SOL: 12.8 ± 5.3 vs. 13.2 ± 5.3 μV, p = 0.453; MG: 8.9 ± 4.2 vs. 9.9 ± 5.3 μV, p = 0.313). Likewise, the number of identified motor units did not differ between tasks for the SOL (8.3 ± 4.5 vs. 8.8 ± 5.3, p = 0.568) or the MG (13.2 ± 4.5 vs. 15.2 ± 9.3, p = 0.185), and firing rates were similar across tasks in both muscles (SOL: 9.9 ± 1.6 vs. 9.8 ± 1.7 pps, p = 0.907; MG: 10.4 ± 0.8 vs. 10.2 ± 1.3 pps, p = 0.515). In contrast, the CoV of low-frequency EMG was significantly lower during standing than during isometric voluntary contraction in both muscles (SOL: p < 0.001; MG: p < 0.001). Moreover, during standing, the MG exhibited a significantly higher CoV than the SOL (p < 0.001) (Fig. 4). On the other hand, no significant difference was observed between the SOL and MG during isometric voluntary contraction (p = 0.852).

**Figure 3.**
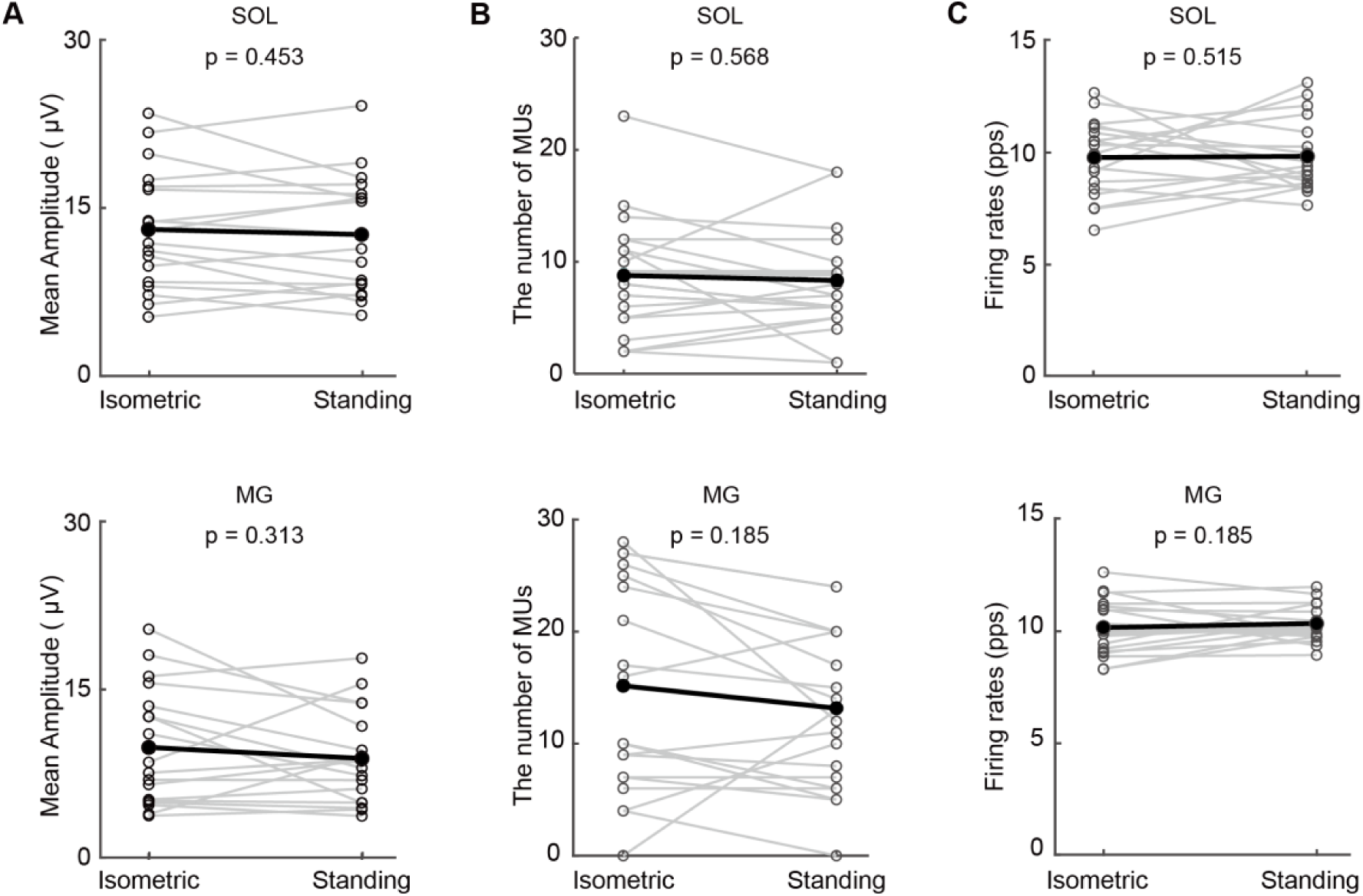
A. Mean EMG amplitude for the isometric voluntary contraction and standing tasks. B. Number of motor units (MUs) for the two tasks. C. Mean firing rates for the two tasks.

**Figure 4.**
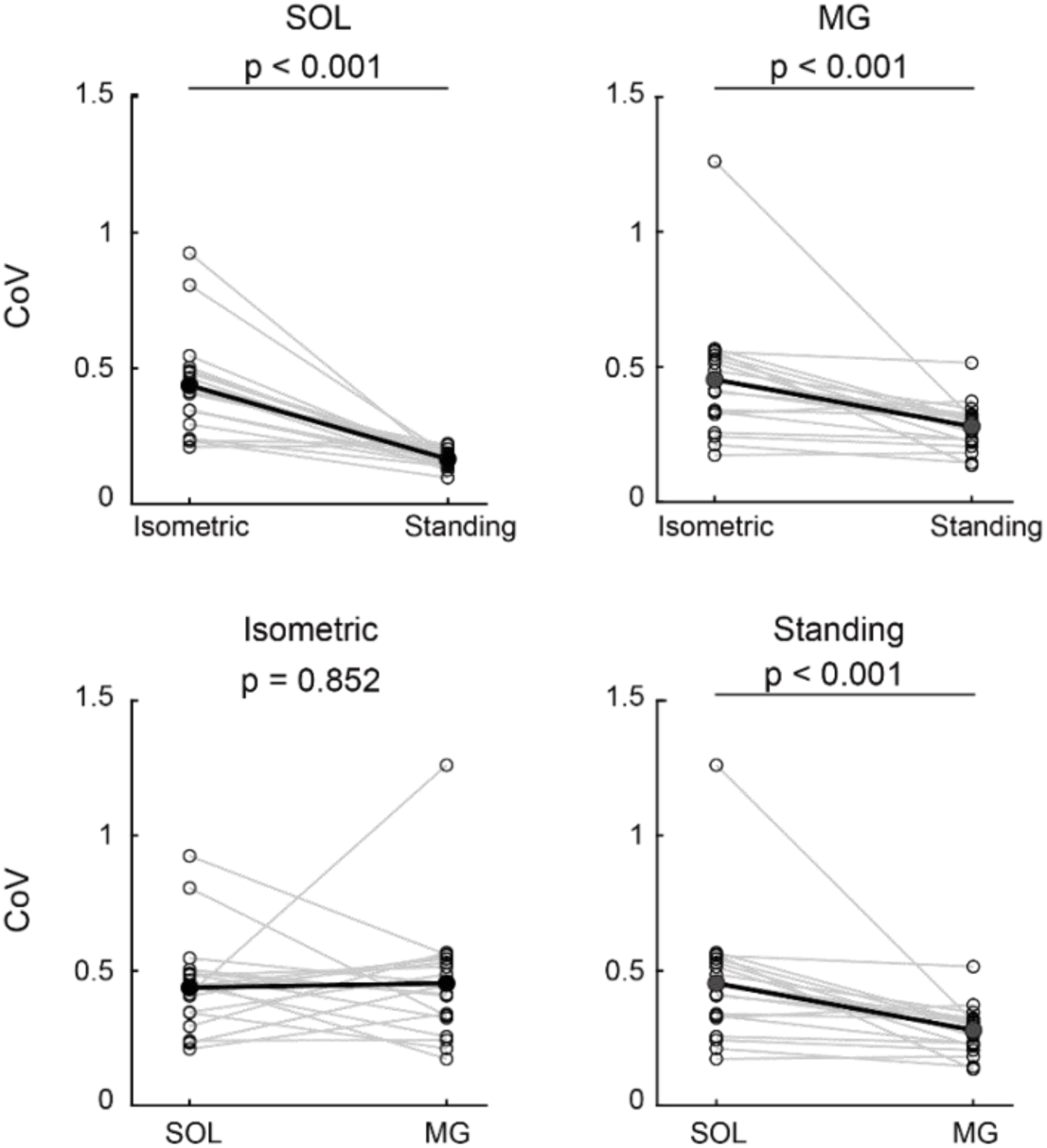
Comparison of the CoV of low-frequency EMG between the isometric voluntary contraction task and standing, and between SOL and MG.

### PSD of surface EMG

We examined the PSD of surface EMG. Figure 5A shows the averaged PSD across participants for four conditions. The PSD of the MG during standing showed higher PSD in the delta and alpha bands than the SOL. The average PSD in each frequency band is shown in Figure 5B.

**Figure 5.**
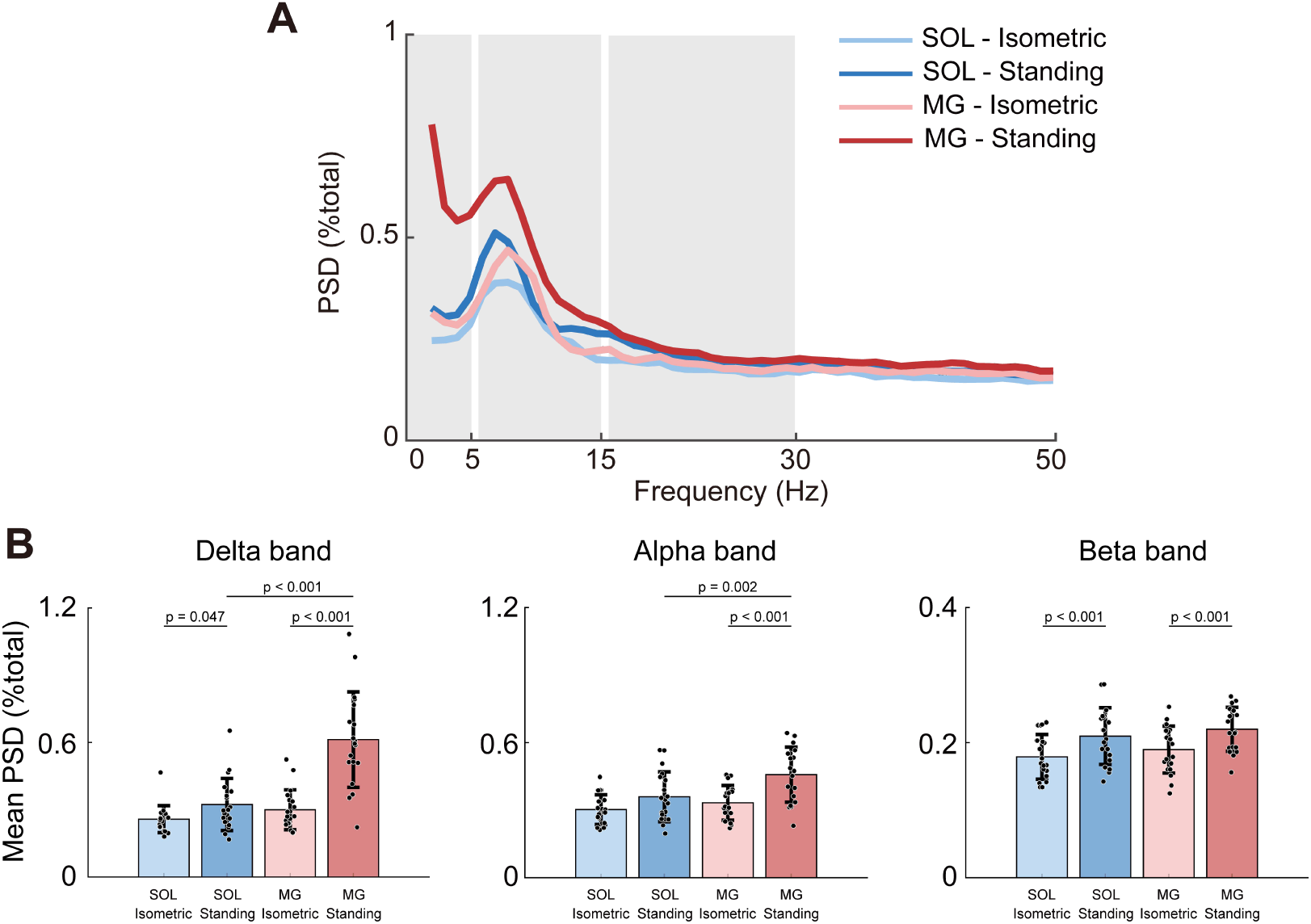
PSD of surface EMG. A. PSD of the MG and SOL during the isometric voluntary contraction task and the standing task. Averaged PSD across participants. Gray shaded areas correspond to frequency bands used for comparisons among the conditions. B. Comparisons of PSD between tasks (standing and isometric voluntary contraction) and between muscles (MG and SOL) across the three frequency bands. Bars represent the mean, error bars indicate the standard deviation, and dots represent individual data points.

In the delta band, we observed a significant main effect of posture on the PSD (F (1, 19) = 39.170, p < 0.001, η^2^ = 0.349) and a significant main effect of muscle (F (1, 19) = 48.160, p < 0.001, η^2^ = 0.291). Moreover, a significant interaction between posture and muscle was demonstrated (F (1, 19) = 35.580, p < 0.001, η^2^ = 0.187). Post-hoc analyses revealed that in the SOL, standing showed significantly greater delta-band PSD than voluntary contraction (p = 0.047, d = 0.610). In the MG, standing also showed greater delta-band PSD than voluntary contraction (p < 0.001, d = 1.570). In addition, during standing, the MG exhibited significantly greater delta-band PSD than the SOL (p < 0.001, d = 1.650). No significant differences were observed between SOL and MG in the voluntary contraction condition, although a moderate effect size was present (p = 0.091, d = 0.490).

In the alpha band, we observed a significant main effect of posture on the PSD (F (1, 19) = 21.240, p < 0.001, η^2^ = 0.540) and a significant main effect of muscle (F (1, 19) = 16.500, p < 0.001, η^2^ = 0.480). Moreover, a significant interaction between posture and muscle was demonstrated (F (1, 19) = 4.570, p = 0.046, η^2^ = 0.200). Post-hoc analyses revealed that in the MG, standing also showed greater alpha-band PSD than voluntary contraction (p < 0.001, d = 1.110). In addition, during standing, the MG exhibited significantly greater alpha-band PSD than the SOL (p = 0.002, d = 0.980). In the SOL, there was no significant difference between standing and voluntary contraction (p = 0.105, d = 0.523). Furthermore, no significant differences were observed between SOL and MG in the voluntary contraction task (p = 0.376, d = 0.310).

In the beta band, we observed a significant main effect of posture on the PSD (F (1, 19) = 25.360, p < 0.001, η^2^ = 0.580) and a significant main effect of muscle (F (1, 19) = 2.820, p = 0.110, η^2^ = 0.140). No significant interaction between posture and muscle was demonstrated (F (1, 19) < 0.001, p = 0.956, η^2^ < 0.001). Post-hoc analyses revealed that in the MG, PSD was significantly greater during standing than voluntary contraction (p < 0.001, d = 1.13). Similarly, in the SOL, PSD during standing was significantly larger than during voluntary contraction (p < 0.001, d = 0.70). On the other hand, no significant difference between the MG and SOL was observed in either the standing or voluntary contraction (standing: p = 0.368, d = 0.320; SOL: p = 0.368, d = 0.260).

### PSD of EEG

We examined the PSD of the EEG to compare the cortical activity between the standing and isometric voluntary contraction tasks. Figure 6A shows the averaged PSD across participants for three conditions. The average PSD in each frequency band is shown in Figure 6B.

**Figure 6.**
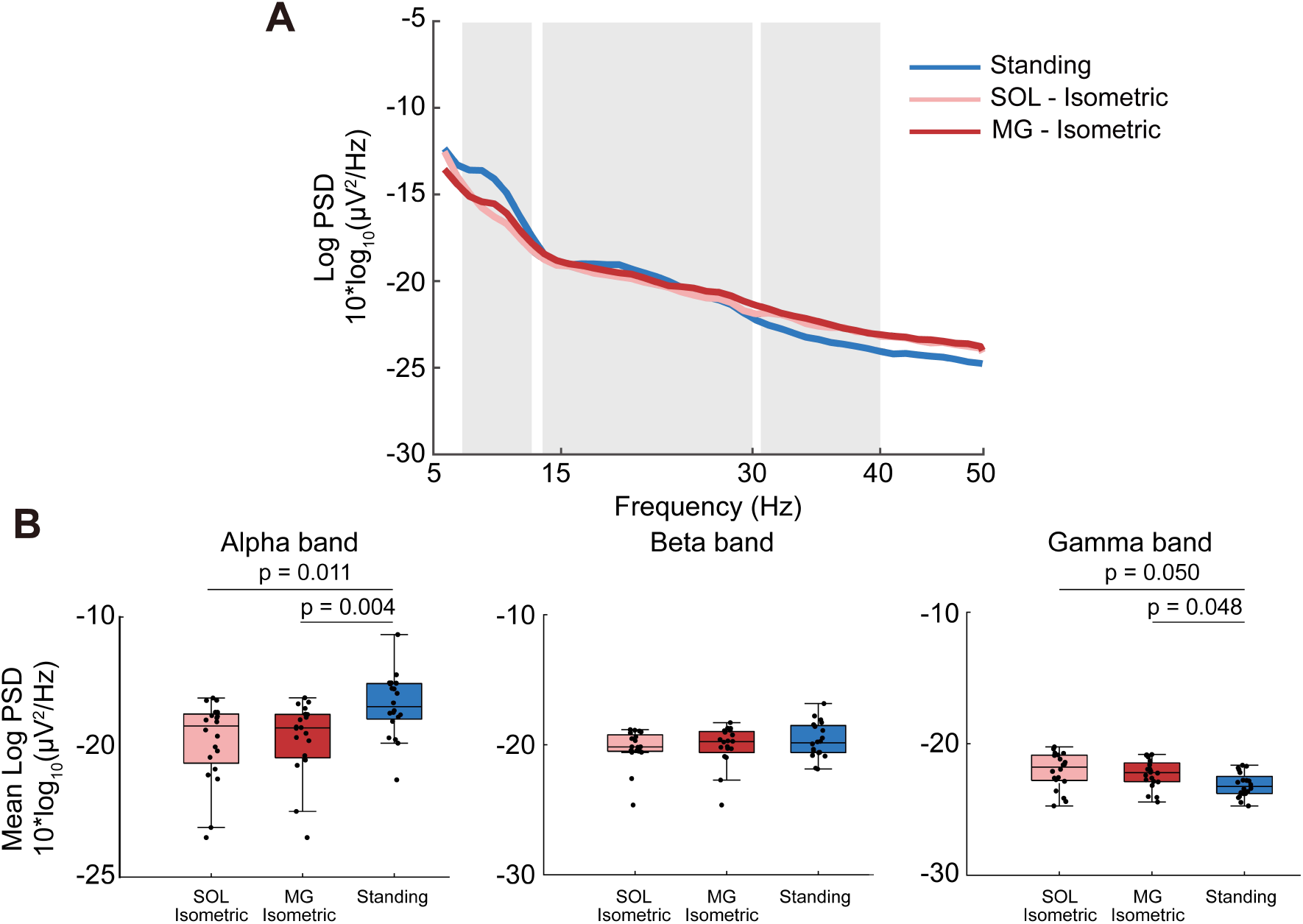
PSD of the sensorimotor area. **A**. PSD of EEG during the isometric voluntary contraction task and the standing task. Averaged PSD across participants. Gray shaded areas correspond to frequency bands used for comparisons among the conditions. **B**. Boxplots of mean log-transformed PSD in the alpha, beta, and gamma bands for each condition (SOL–isometric, MG–isometric, and standing). The horizontal lines within the boxes represent the **median**, while the bars and error bars indicate the mean and standard deviation, respectively. Individual data points are represented by dots.

In the alpha band PSD, the Friedman test revealed a significant main effect of condition (χ^2^ (2) = 9.100, p = 0.011, W = 0.240). Post-hoc analyses showed that standing exhibited significantly higher PSD than both the SOL and MG isometric contractions (standing vs. SOL: p = 0.011, r = 0.670; standing vs. MG: p = 0.004, r = 0.700). No significant difference was observed between SOL and MG during the isometric voluntary contraction tasks (p = 1.000, r = 0.040).

In the beta band PSD, the main effect of condition was not significant (χ^2^ (2) = 0.080, p = 0.960, W < 0.010).

In the gamma band PSD, condition had a significant effect (χ^2^ (2) = 7.320, p = 0.026, W = 0.190). Post-hoc analysis indicated that standing showed lower gamma-band PSD than the MG and SOL isometric voluntary contraction tasks (standing vs. MG: p = 0.048, r = 0.540; standing vs. SOL: p = 0.050, r = 0.560). No significant difference was observed between SOL and MG during isometric voluntary contraction (p = 1.000, r = 0.060).

### Intramuscular coherence (Fig.7)

We examined the synchronization inputs to the MUs by using IMC analysis.

**Figure 7.**
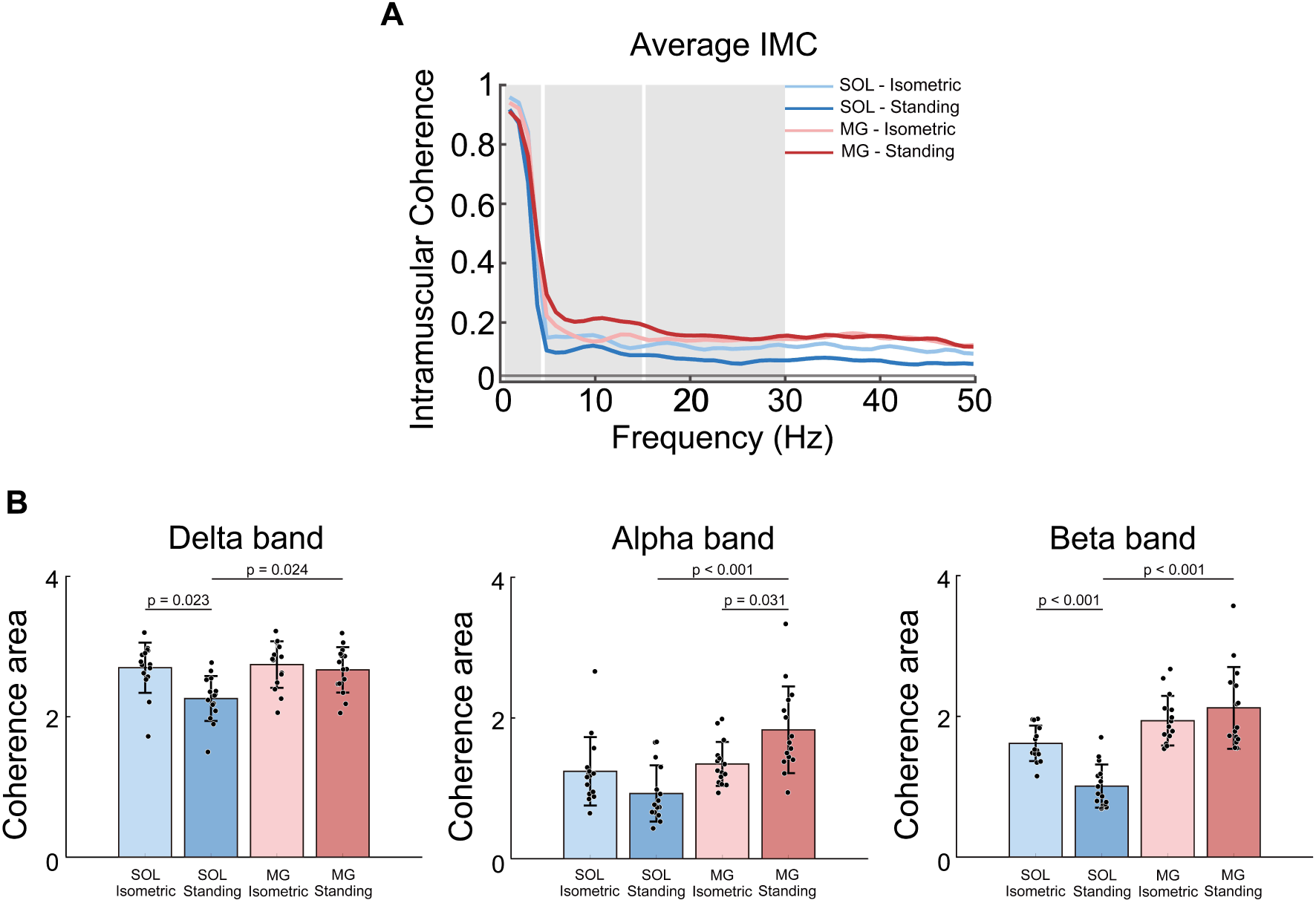
Coherence of MU activities. A. IMC of the MG and SOL during the isometric voluntary contraction task and the standing task. Averaged IMC across participants. Gray horizontal lines indicate a significant threshold for coherence value. Gray shaded areas correspond to frequency bands used for comparisons among the conditions. B. Comparisons of coherence values between tasks (standing vs. isometric voluntary contraction) and between muscles (MG vs. SOL) across the three frequency bands. Bars represent the mean, error bars indicate the standard deviation, and dots represent individual data points.

Figure 7A shows the averaged IMC across participants for four conditions. In all conditions, coherence values exceeded the significance level (1-50 Hz). Overall, the MG tended to show higher coherence values than the SOL across the entire frequency range during the standing task, whereas in the isometric voluntary contraction task, there was only a slight difference between the muscles. The IMC areas in each frequency band are shown in Figure 7B.

In the delta band, we observed a significant main effect of posture on the coherence area (F = 8.067, p = 0.013, η^2^ = 0.117) and a significant main effect of muscle (F (1, 14) = 9.429, p = 0.008, η^2^ = 0.091). Moreover, a significant interaction between posture and muscle was demonstrated (F (1, 14) = 5.118, p = 0.040, η^2^ = 0.058). Post-hoc analyses revealed that in the SOL, standing showed significantly smaller IMC than voluntary contraction (p = 0.023, d = 0.840). In addition, during standing, the SOL exhibited significantly smaller IMC than the MG (p = 0.024, d = 1.14). No significant differences were observed between SOL and MG in the voluntary contraction (p = 1.000, d = 0.090), nor between voluntary contraction and standing in MG (p = 1.000, d = 0.190).

In the alpha band, we observed a significant main effect of muscle on the coherence area (F (1, 14) = 28.100, p < 0.001, η^2^ = 0.206) and a significant interaction between posture and muscle (F (1, 14) = 39.890, p < 0.001, η^2^ = 0.129). No significant main effect of posture was observed (F (1, 14) = 0.493, p = 0.494, η^2^ = 0.006). Post hoc analyses revealed that in the MG, standing showed significantly larger IMC than voluntary contraction (p = 0.031, d = 0.800). In addition, during standing, the MG exhibited significantly larger IMC than the SOL (p < 0.001, d = 2.430). No significant difference was observed between voluntary contraction and standing in the SOL (p = 0.069, d = 0.70), nor between the SOL and the MG in the voluntary contraction (p = 1.000, d = 0.210).

In the beta band, we observed a significant main effect of muscle on the coherence area (F (1, 14) = 56.220, p < 0.001, η^2^ = 0.397), and a significant interaction between posture and muscle (F (1, 14) = 38.300, p < 0.001, η^2^ = 0.121). No significant main effect of posture was observed (F (1, 14) = 4.091, p = 0.063, η^2^ = 0.035). Post hoc analyses revealed that in the SOL, standing showed significantly smaller IMC than voluntary contraction (p < 0.001, d = 1.330). Moreover, during standing, the MG exhibited significantly larger IMC than the SOL (p < 0.001, d = 3.060). No significant difference was observed between voluntary contraction and standing in the MG (p = 0.704, d = 0.370), nor between the SOL and MG in the voluntary contraction (p = 0.118, d = 0.630).

### Corticomuscular coherence (Fig.8)

The connectivity between the cortex and spinal motor neurons was assessed using CMC analysis. Figure 8A shows the averaged CMC across participants for the four conditions. Overall, in contrast to IMC, no clear differences between the muscles were observed in either the standing or the isometric voluntary contraction task. With respect to task differences, CMC tended to be lower during standing than during the isometric voluntary contraction task in both muscles. The CMC areas in each frequency band are shown in Figure 8B.

**Figure 8.**
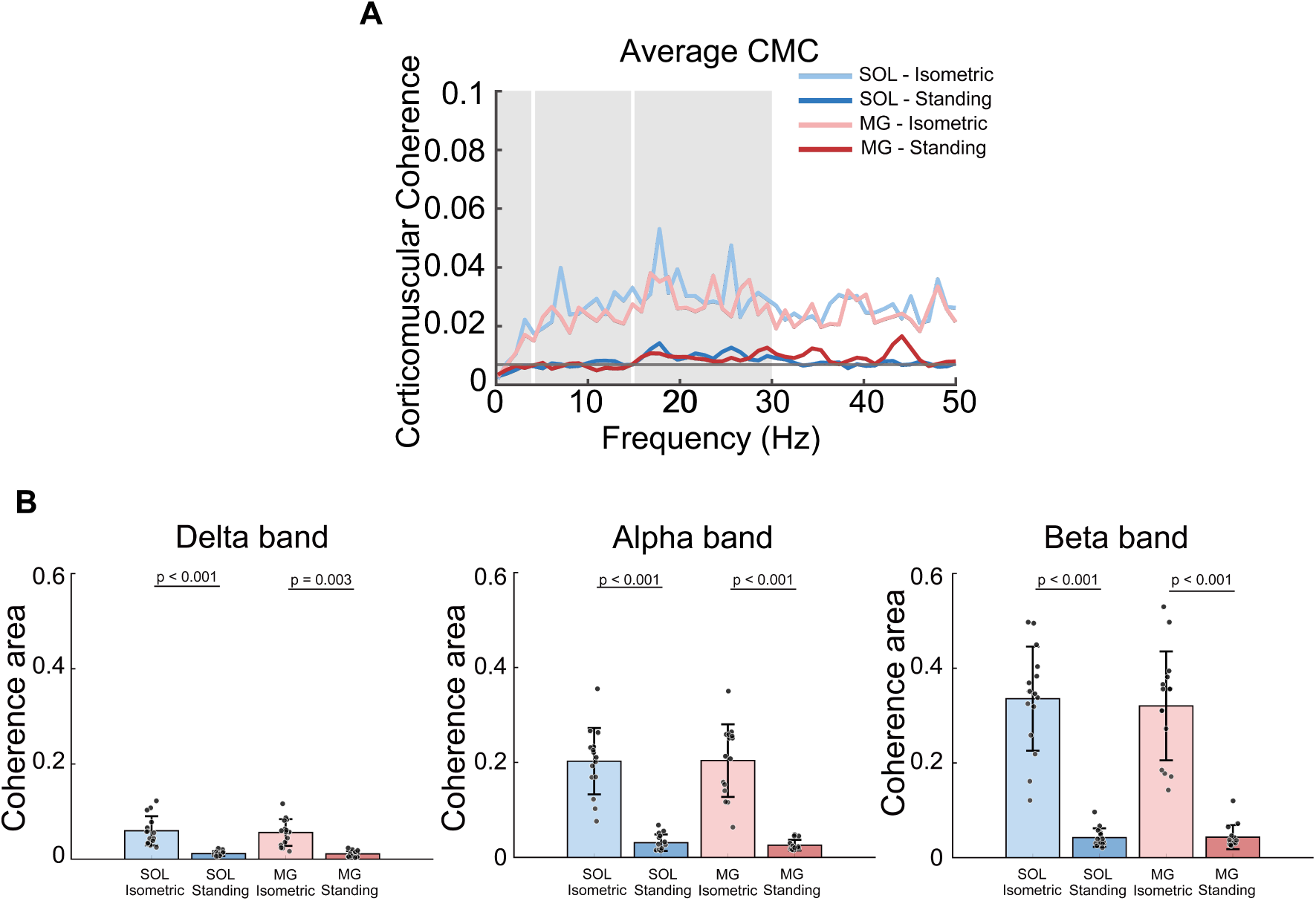
Coherence between EEG and MU activities. A. CMC of the MG and SOL during the isometric voluntary contraction task and the standing task. Averaged CMC across participants. Gray horizontal lines indicate a significant threshold for coherence value. Gray shaded areas correspond to frequency bands used for comparisons among the conditions. B. Comparisons of coherence values between tasks (standing vs. isometric voluntary contraction) and between muscles (MG vs. SOL) across the three frequency bands. Bars represent the mean, error bars indicate the standard deviation, and dots represent individual data points.

In the delta band, we observed a significant main effect of posture on the coherence area (F (1, 14) = 41.150, p < 0.001, η^2^ = 0.505). No significant main effect of muscle (F (1, 14) = 1.797, p = 0.201, η^2^ = 0.012) nor a significant interaction between posture and muscle (F (1, 14) = 1.935, p = 0.186, η^2^ = 0.012) was observed. Post hoc paired t-tests with Bonferroni correction revealed that both in the SOL and MG, voluntary contraction showed significantly larger CMC than standing (SOL: p < 0.001, d = 1.600; MG: p = 0.003, d = 1.110). No significant difference between muscles in either posture (isometric voluntary contraction: p = 0.660, d = 0.380; standing: p = 1.000, d = 0.001).

In the alpha band, we observed a significant main effect of posture on the coherence area (F (1, 14) = 96.980, p < 0.001, η^2^ = 0.661). No significant main effect of muscle (F (1, 14) = 0.807, p = 0.384, η^2^ = 0.004), nor a significant interaction between posture and muscle (F (1, 14) = 0.588, p = 0.460, η^2^ = 0.003). Post hoc paired t-tests with Bonferroni correction revealed that both in the SOL and MG, voluntary contraction showed significantly larger CMC than standing (SOL: p < 0.001, d = 2.290; MG: p < 0.001, d = 1.670). No significant difference between muscles in either posture (isometric voluntary contraction: p = 1.000, d = 0.230; standing: p = 1.000, d = 0.050).

In the beta band, there was a significant main effect of posture on the coherence area (F (1, 14) = 97.55, p < 0.001, η² = 0.678), as well as a significant interaction between posture and muscle (F (1, 14) = 5.07, p = 0.041, η² = 0.007). No significant main effect of muscle (F (1, 14) = 1.840, p = 0.197, η^2^ = 0.009). Post hoc paired t-tests with Bonferroni correction revealed that both in the SOL and MG, voluntary contraction showed significantly larger CMC than standing (SOL: p < 0.001, d = 2.680; MG: p < 0.001, d = 2.020). No significant difference between muscles in either posture (isometric voluntary contraction: p = 0.317, d = 0.490; standing: p = 1.000, d = 0.050).

## Discussion

Many studies have demonstrated that the MG and SOL play distinct functional roles during quiet standing (Masani *et al*., 2003; Héroux *et al*., 2014). However, the neural mechanisms that give rise to this functional divergence have remained unclear. To address this gap, this study compared the neural control strategies of the SOL and MG during quiet standing, focusing on the synaptic inputs underlying MU behavior.

Our findings reveal clear muscle-dependent differences in the neural inputs to spinal motor neurons, providing the first evidence that the distinct postural functions of the MG and SOL arise from different patterns of neural input at the motor neuron level.

### Neural inputs underlying MG activation during quiet standing

Alpha-band IMC, which reflects Ia afferent and vestibulospinal inputs (Liv Hansen & Bo Nielsen, 2004; Dakin *et al*., 2007), was larger in the MG than in the SOL during standing (Fig. 7). In addition, while alpha-band IMC did not differ between tasks in the SOL, the MG showed a clear increase during standing relative to isometric voluntary contraction. Therefore, these findings indicate that only the MG exhibits a task-dependent increase in alpha-band IMC. Given that the modulation of Ia afferent excitability is largely similar between the SOL and MG during standing (Tokuno *et al*., 2007, 2008), the observed difference in alpha-band IMC is unlikely to arise from Ia afferent input. Instead, it is more plausibly explained by muscle-specific differences in vestibular input. The MG shows larger vestibular-evoked responses than the SOL (Dakin *et al*., 2016), and vestibular input increases during standing when postural demands are greater than in sitting (Forbes *et al*., 2015). To further evaluate the functional significance of the vestibular contribution, we analyzed alpha-band PSD from surface EMG and found that alpha power in the MG was higher during standing than during the isometric voluntary contraction task (Fig. 5). Given that alpha-band PSD during standing contributes to increased ankle stiffness (Kouzaki & Fukunaga, 2008), the elevated alpha power likely reflects enhanced vestibulospinal input that maintains MG tone and ankle stiffness to stabilize postural sway. Therefore, the present findings suggest that vestibulospinal input becomes more strongly engaged during standing and contributes more prominently to MG activation than to SOL activation. This interpretation aligns with the MG’s established role in generating corrective ankle torques to counter postural sway (Masani *et al*., 2003; Héroux *et al*., 2014), providing evidence that the relative contribution of descending inputs differs between the MG and SOL during postural control and clarifying the neural mechanisms that differentiate their functional contributions.

On the other hand, beta-band IMC, which reflects corticospinal and reticulospinal inputs (Thompson *et al*., 2019; Bräcklein *et al*., 2022), demonstrated that the MG exhibited greater IMC than the SOL during standing (Fig. 7). In the task comparison, the SOL showed a clear reduction in beta-band IMC during standing relative to voluntary contraction (Fig. 8). This decrease paralleled a reduction in CMC, which specifically reflects corticospinal connectivity (Conway *et al*., 1995). In contrast, although the MG also showed reduced CMC during standing, its beta-band IMC remained unchanged across tasks. This divergence between IMC and CMC, specifically in the MG, suggests that the preserved beta-band IMC during standing is not due to sustained corticospinal input, but rather to an increased contribution of reticulospinal input—a pathway known to become more engaged when postural demands are elevated. Reticulospinal pathways modulate the level of muscle tone during postural adjustments (Takakusaki, 2017). In addition, high-threshold motor neurons appear to be more strongly influenced by the reticulospinal than the corticospinal tract (Baker, 2011; Glover & Baker, 2020). Because MG MUs have higher recruitment thresholds than those of the SOL (Hali *et al*., 2020), they may be more responsive to reticulospinal input. Moreover, the MG should generate rapid, phasic torque to correct postural sway, requiring dynamically modulated muscle tone—another condition that likely increases reliance on reticulospinal input during standing. To further interpret this pattern, EEG PSD was analyzed from the sensorimotor cortex. Alpha-band PSD was lower during isometric voluntary contraction than standing, consistent with increased cortical processing demands during voluntary force control (Ortiz et al., 2023). Moreover, gamma-band PSD was also higher during isometric voluntary contraction than during standing. Given that higher gamma activity reflects enhanced early stage of cortical processing of somatosensory input (Fukuda *et al*., 2010), the reduced gamma power during standing suggests that rapid sensory–cortical processing demands are lower in standing than in the isometric voluntary contraction task. In contrast, beta-band PSD did not differ between tasks, indicating a comparable level of tonic motor output across tasks (Engel & Fries, 2010). These EEG findings support the interpretation that cortical drive does not substantially increase during standing, and thus cannot account for the maintained beta-band IMC in the MG. Therefore, the present findings indicate that, compared with the SOL, the MG receives stronger reticulospinal input during standing. This enhanced subcortical input likely enables the MG to rapidly adjust muscle tone and generate phasic torque needed to counter postural sway.

### Neural mechanisms supporting continuous SOL activation during standing

Delta-band IMC, which is associated with fluctuations in muscle force (McManus *et al*., 2016), was smaller in the SOL than in the MG (Fig. 7). Moreover, while delta-band IMC remained unchanged across tasks in the MG, the SOL showed a clear reduction during standing compared with isometric voluntary contraction. These findings suggest that the SOL produces steadier force, whereas the MG generates more variable force. Consistent with this interpretation, the low-frequency EMG analysis revealed a smaller CoV in the SOL than in the MG (Fig. 4), and EMG PSD analysis also showed greater delta-band power in the MG than in the SOL (Fig.5). Taken together, these results confirm the previously described functional distinction between the two muscles: the MG engages in phasic activity aligned with postural sway, whereas the SOL provides tonic activity to support body weight (Héroux *et al*., 2014).

On the other hand, overall IMC also differed between the two muscles. As shown in Fig. 7, IMC was smaller in the SOL than in the MG across all bands. This pattern suggests that the SOL receives a weaker common neural input and maintains characteristically low MU synchrony during standing. Such low synchrony is well suited to the SOL’s tonic, steady torque production, which is essential for supporting body weight during standing. The degree of MU synchrony influences muscle twitch force: high MU synchrony enhances the efficiency of twitch force generation, whereas low synchrony ensures more stable force output (Yao *et al*., 2000; Taylor *et al*., 2002). Consequently, the low MU synchrony observed in the SOL makes it less suited for generating rapid, twitch-like force responses to postural perturbations, but it provides a highly stable force-generation strategy that enables continuous, low-variability plantarflexion torque during quiet standing. In addition, the SOL can sustain such continuous activity even with relatively small synaptic input because its motor units have lower recruitment thresholds than those of the MG (Hali *et al*., 2020). This property allows the SOL to maintain firing without requiring strong common input. Furthermore, because slow-twitch–dominant muscles exhibit longer-lasting persistent inward currents (PICs) than fast-twitch–dominant muscles (Heckman *et al*., 2008), the SOL may amplify and sustain descending and afferent inputs more effectively than the MG. Such intrinsic motor neuron properties likely support the SOL’s continuous, tonic activation during standing.

### Distinct neural control strategies of the SOL and MG during quiet standing (Fig.9)

In summary, our results revealed that the neural mechanisms controlling the SOL and MG during postural control are distinct. These findings offer new insights into the neural mechanisms underlying the long-described functional distinction between the MG and SOL during standing. The MG should generate rapid, phasic plantarflexion torque to counter postural sway, and our results suggest that this role is supported by a high-gain neural control strategy that relies on strong vestibulospinal and reticulospinal inputs. In contrast, the SOL should continuously generate steady plantarflexion torque to support body weight. Its neural strategy relies on relatively weak common synaptic input and low MU synchrony, supporting stable, low-variability plantarflexion torque that is well suited for tonic antigravity function. Therefore, these findings elucidate the neural mechanisms that underlie the distinct functional contributions of the SOL and MG during standing.

**Figure 9.**
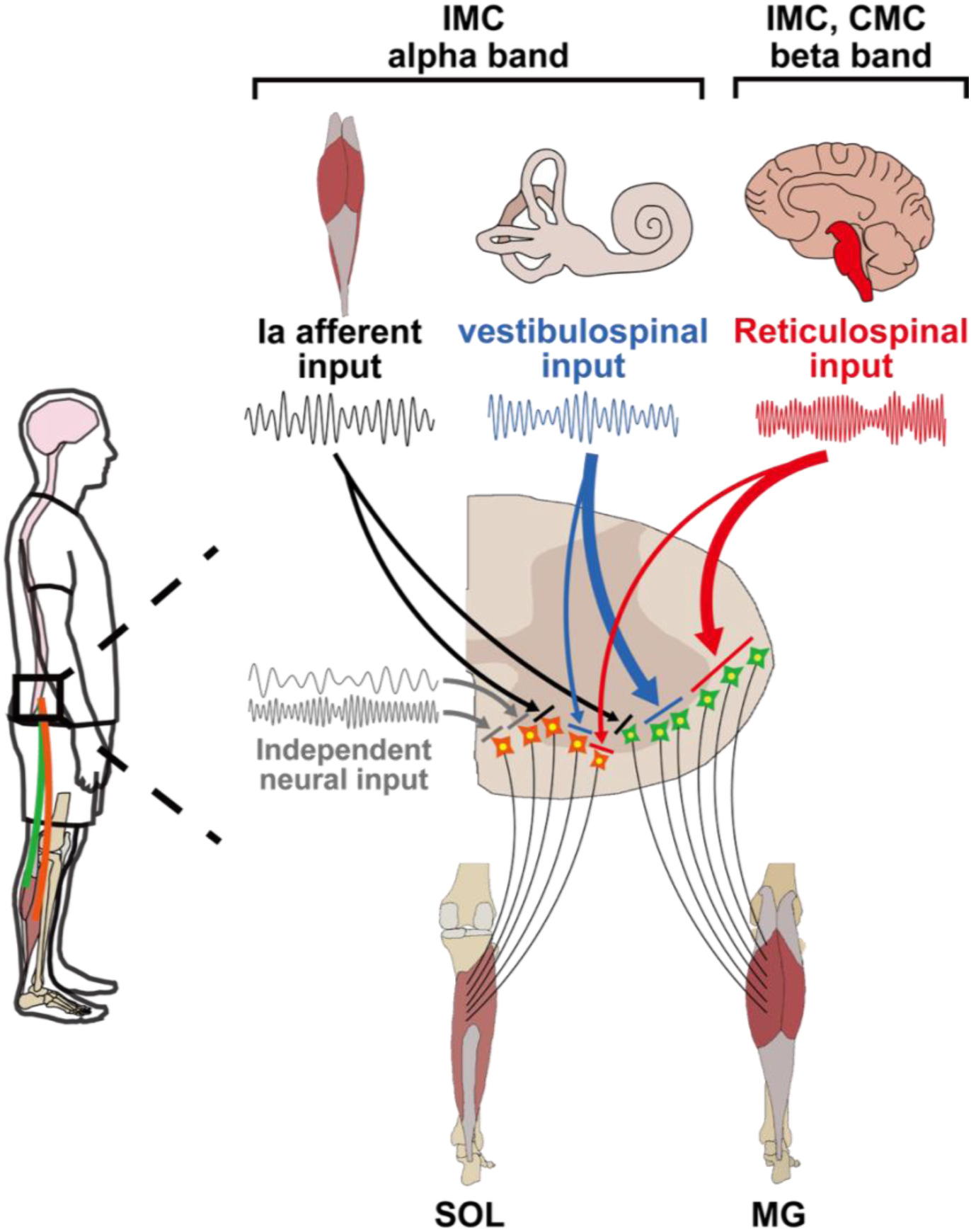
Schematic illustration showing neural control mechanisms for the MG and SOL during standing. Based on the IMC and CMC results, the MG appears to be primarily controlled by vestibulospinal and reticulospinal inputs. In contrast, the SOL receives less common synaptic input, suggesting a neural control strategy characterized by reduced synchrony among motor neurons.

### Limitation

A limitation of this study is that we did not measure kinetics using motion capture. Therefore, it was not possible to distinguish whether the MUs identified in the MG were recruited to generate knee flexion torque or ankle plantarflexion torque. However, during quiet standing, postural control primarily depends on ankle plantarflexion torque, with only a minor contribution from the knee joint (Winter, 1995).

In addition, the isometric voluntary contraction task was controlled based on muscle activity rather than ankle joint torque in this study. Thus, the actual torque exerted at the ankle joint may not have been identical across tasks. However, controlling by joint torque would not have allowed us to directly compare standing with the isometric voluntary contraction task for each muscle (SOL and MG), which was the primary aim of this study. For this reason, muscle activity was used as the feedback variable.

## Conclusion

Our study explored the differential neural control mechanisms of the SOL and MG during quiet standing, revealing that these muscles rely on distinct patterns of neural input. We observed that the MG exhibited larger IMC than the SOL, particularly in the alpha and beta bands. Importantly, our findings suggest that the MG relies more heavily on neural inputs, likely from vestibulospinal and reticulospinal pathways, to generate phasic ankle plantarflexion torque for postural sway control. In contrast, the SOL depends on more dispersed inputs that support continuous and stable force production. Therefore, the current study offers valuable insights into the neuromechanical basis of postural control, thereby advancing our understanding of how the central nervous system coordinates synergistic muscles to maintain standing.

## Additional information

The data that support the findings of this study are available on request from the corresponding author.

## Author contribution

H.A., N.K., H.Y., and K.N. conceptualized and designed the research. H.A. and N.H. performed the experiments. H.A. analyzed the data. All authors interpreted the results of the experiments. H.A. drafted the manuscript. N.K., N.H., H.Y., and K.N. critically revised the manuscript for important intellectual content. All authors have read and approved the final version of this manuscript and agree to be accountable for all aspects of the work in ensuring that questions related to the accuracy or integrity of any part of the work are appropriately investigated and resolved. All persons designated as authors qualify for authorship, and all those who qualify for authorship are listed.

## Reference

1. Baker SN (2011). The primate reticulospinal tract, hand function and functional recovery. The Journal of Physiology 589, 5603–5612.

2. Bräcklein M, Barsakcioglu DY, Del Vecchio A, Ibáñez J & Farina D (2022). Reading and Modulating Cortical β Bursts from Motor Unit Spiking Activity. J Neurosci 42, 3611–3621.

3. Buchthal F & Schmalbruch H (1980). Motor unit of mammalian muscle. Physiological Reviews 60, 90–142.

4. Cabral HV, Cudicio A, Bonardi A, Del Vecchio A, Falciati L, Orizio C, Martinez-Valdes E & Negro F (2024a). Neural Filtering of Physiological Tremor Oscillations to Spinal Motor Neurons Mediates Short-Term Acquisition of a Skill Learning Task. eNeuro 11, ENEURO.0043-24.2024.

5. Cabral HV, Inglis JG, Cudicio A, Cogliati M, Orizio C, Yavuz US & Negro F (2024b). Muscle contractile properties directly influence shared synaptic inputs to spinal motor neurons. The Journal of Physiology 602, 2855–2872.

6. Cadzow J & Solomon O (1987). Linear modeling and the coherence function. *IEEE Trans Acoust, Speech*, Signal Process 35, 19–28.

7. Castronovo AM, Negro F & Farina D (2015). Theoretical Model and Experimental Validation of the estimated proportions of common and independent input to motor neurons. In 2015 37th Annual International Conference of the IEEE Engineering in Medicine and Biology Society (EMBC), pp. 254–257. IEEE, Milan. Available at: http://ieeexplore.ieee.org/document/7318348/ [Accessed September 2, 2025].

8. Cohen J (2009). Statistical power analysis for the behavioral sciences, 2. ed., reprint. Psychology Press, New York, NY.

9. Contreras-Hernandez I, Falla D, Arvanitidis M, Negro F, Jimenez-Grande D & Martinez-Valdes E (2025). Load and muscle-dependent changes in triceps surae motor unit firing properties in individuals with non-insertional Achilles tendinopathy. The Journal of PhysiologyJP287588.

10. Conway BA, Halliday DM, Farmer SF, Shahani U, Maas P, Weir AI & Rosenberg JR (1995). Synchronization between motor cortex and spinal motoneuronal pool during the performance of a maintained motor task in man. The Journal of Physiology 489, 917–924.

11. Dakin CJ, Héroux ME, Luu BL, Inglis JT & Blouin J-S (2016). Vestibular contribution to balance control in the medial gastrocnemius and soleus. Journal of Neurophysiology 115, 1289–1297.

12. Dakin CJ, Son GML, Inglis JT & Blouin J (2007). Frequency response of human vestibular reflexes characterized by stochastic stimuli. The Journal of Physiology 583, 1117–1127.

13. Del Vecchio A, Casolo A, Negro F, Scorcelletti M, Bazzucchi I, Enoka R, Felici F & Farina D (2019). The increase in muscle force after 4 weeks of strength training is mediated by adaptations in motor unit recruitment and rate coding. The Journal of Physiology 597, 1873–1887.

14. Edelman BJ, Baxter B & He B (2016). EEG Source Imaging Enhances the Decoding of Complex Right-Hand Motor Imagery Tasks. IEEE Trans Biomed Eng 63, 4–14.

15. Farmer SF, Gibbs J, Halliday DM, Harrison LM, James LM, Mayston MJ & Stephens JA (2007). Changes in EMG coherence between long and short thumb abductor muscles during human development. The Journal of Physiology 579, 389–402.

16. Forbes PA, Siegmund GP, Schouten AC & Blouin J-S (2015). Task, muscle and frequency dependent vestibular control of posture. Front Integr Neurosci; DOI: 10.3389/fnint.2014.00094.

17. Fukuda M, Juhász C, Hoechstetter K, Sood S & Asano E (2010). Somatosensory-related gamma-, beta- and alpha-augmentation precedes alpha- and beta-attenuation in humans. Clinical Neurophysiology 121, 366–375.

18. Glover IS & Baker SN (2020). Cortical, Corticospinal, and Reticulospinal Contributions to Strength Training. J Neurosci 40, 5820–5832.

19. Gomes MM, Jenz ST, Beauchamp JA, Negro F, Heckman CJ & Pearcey GEP (2024). Voluntary co-contraction of ankle muscles alters motor unit discharge characteristics and reduces estimates of persistent inward currents. The Journal of Physiology 602, 4237–4250.

20. Hali K, Dalton BH, Harwood B, Fessler AF, Power GA & Rice CL (2020). Differential Modulation of Motor Unit Properties from the Separate Components of the Triceps Surae in Humans. Neuroscience 428, 192–198.

21. Hayashi R, Tako K, Tokuda T & Yanagisawa N (1992). Comparison of amplitude of human soleus H-reflex during sitting and standing. Neuroscience Research 13, 227–233.

22. Heckman CJ, Johnson M, Mottram C & Schuster J (2008). Persistent Inward Currents in Spinal Motoneurons and Their Influence on Human Motoneuron Firing Patterns. Neuroscientist 14, 264–275.

23. Héroux ME, Dakin CJ, Luu BL, Inglis JT & Blouin J-S (2014). Absence of lateral gastrocnemius activity and differential motor unit behavior in soleus and medial gastrocnemius during standing balance. Journal of Applied Physiology 116, 140–148.

24. Jenz ST, Beauchamp JA, Gomes MM, Negro F, Heckman CJ & Pearcey GEP (2023). Estimates of persistent inward currents in lower limb motoneurons are larger in females than in males. Journal of Neurophysiology 129, 1322–1333.

25. Kayser J & Tenke CE (2006). Principal components analysis of Laplacian waveforms as a generic method for identifying ERP generator patterns: I. Evaluation with auditory oddball tasks. Clinical Neurophysiology 117, 348–368.

26. Kouzaki M & Fukunaga T (2008). Frequency features of mechanomyographic signals of human soleus muscle during quiet standing. Journal of Neuroscience Methods 173, 241–248.

27. Liv Hansen N & Bo Nielsen J (2004). The effect of transcranial magnetic stimulation and peripheral nerve stimulation on corticomuscular coherence in humans. The Journal of Physiology 561, 295–306.

28. Loram ID & Lakie M (2002). Direct measurement of human ankle stiffness during quiet standing: the intrinsic mechanical stiffness is insufficient for stability. The Journal of Physiology 545, 1041–1053.

29. Loram ID, Maganaris CN & Lakie M (2005). Human postural sway results from frequent, ballistic bias impulses by soleus and gastrocnemius. The Journal of Physiology 564, 295–311.

30. Makihara Y, Segal RL, Wolpaw JR & Thompson AK (2012). H-reflex modulation in the human medial and lateral gastrocnemii during standing and walking. Muscle and Nerve 45, 116–125.

31. Martinez-Valdes E, Negro F, Falla D, Dideriksen JL, Heckman CJ & Farina D (2020). Inability to increase the neural drive to muscle is associated with task failure during submaximal contractions. Journal of Neurophysiology 124, 1110–1121.

32. Masani K, Popovic MR, Nakazawa K, Kouzaki M & Nozaki D (2003). Importance of Body Sway Velocity Information in Controlling Ankle Extensor Activities During Quiet Stance. Journal of Neurophysiology 90, 3774–3782.

33. McManus L, Hu X, Rymer WZ, Suresh NL & Lowery MM (2016). Muscle fatigue increases beta-band coherence between the firing times of simultaneously active motor units in the first dorsal interosseous muscle. Journal of Neurophysiology 115, 2830–2839.

34. Nazarpour K, Sanei S, Shoker L & Chambers JA (2006). PARALLEL SPACE-TIME-FREQUENCY DECOMPOSITION OF EEG SIGNALS FOR BRAIN COMPUTER INTERFACING.

35. Negro F, Muceli S, Castronovo AM, Holobar A & Farina D (2016). Multi-channel intramuscular and surface EMG decomposition by convolutive blind source separation. J Neural Eng 13, 026027.

36. Pernet C, Garrido MI, Gramfort A, Maurits N, Michel CM, Pang E, Salmelin R, Schoffelen JM, Valdes-Sosa PA & Puce A (2020). Issues and recommendations from the OHBM COBIDAS MEEG committee for reproducible EEG and MEG research. Nat Neurosci 23, 1473–1483.

37. Rossato J, Tucker K, Avrillon S, Lacourpaille L, Holobar A & Hug F (2022). Less common synaptic input between muscles from the same group allows for more flexible coordination strategies during a fatiguing task. Journal of Neurophysiology 127, 421–433.

38. Studnicki A, Seidler RD & Ferris DP (2023). A table tennis serve versus rally hit elicits differential hemispheric electrocortical power fluctuations. Journal of Neurophysiology 130, 1444–1456.

39. Takakusaki K (2017). Functional Neuroanatomy for Posture and Gait Control. JMD 10, 1–17.

40. Taylor AM, Steege JW & Enoka RM (2002). Motor-Unit Synchronization Alters Spike-Triggered Average Force in Simulated Contractions. Journal of Neurophysiology 88, 265–276.

41. Taylor CA, Kopicko BH, Negro F & Thompson CK (2022). Sex differences in the detection of motor unit action potentials identified using high-density surface electromyography. Journal of Electromyography and Kinesiology 65, 102675.

42. Thompson CK, Johnson MD, Negro F, Mcpherson LM, Farina D & Heckman CJ (2019). Exogenous neuromodulation of spinal neurons induces beta-band coherence during self-sustained discharge of hind limb motor unit populations. Journal of Applied Physiology 127, 1034–1041.

43. Tokuno CD, Carpenter MG, Thorstensson A, Garland SJ & Cresswell AG (2007). Control of the triceps surae during the postural sway of quiet standing. Acta Physiologica 191, 229–236.

44. Tokuno CD, Garland SJ, Carpenter MG, Thorstensson A & Cresswell AG (2008). Sway-dependent modulation of the triceps surae H-reflex during standing. Journal of Applied Physiology 104, 1359–1365.

45. Winter D (1995). Human balance and posture control during standing and walking. Gait & Posture 3, 193–214.

46. Winter DA (2009). Biomechanics and motor control of human movement, 4th ed. Wiley, Hoboken, N.J.

47. Yao W, Fuglevand RJ & Enoka RM (2000). Motor-Unit Synchronization Increases EMG Amplitude and Decreases Force Steadiness of Simulated Contractions. Journal of Neurophysiology 83, 441–452.

48. Yokoyama H, Kaneko N, Sasaki A, Saito A & Nakazawa K (2022). Firing behavior of single motor units of the tibialis anterior in human walking as non-invasively revealed by HDsEMG decomposition. J Neural Eng 19, 066033.

